# Gut microbial communities associated with phenotypically divergent populations of the striped stem borer *Chilo suppressalis*

**DOI:** 10.1101/2020.08.10.243956

**Authors:** Haiying Zhong, Jianming Chen, Juefeng Zhang, Fang Li

**Affiliations:** Institute of Plant Protection and Microbiology, Zhejiang Academy of Agricultural Sciences; State Key Laboratory for Managing Biotic and Chemical Threats to the Quality and Safety of Agro-products, Hangzhou 310021, China

**Keywords:** Lepidoptera, midgut, hindgut, microbiota, diet

## Abstract

*Chilo suppressalis* is a serious stem borer of rice and water-oat, however, little is known about the effect of diet and gut compartments on the gut microbial communities of this species. We analyzed the microbial communities in phenotypically divergent populations of *C. suppressalis*. In original and cross-rearing populations, the most dominant phyla were Proteobacteria (16.0% to 96.4%) and Firmicutes (2.3% to 78.9%); the most abundant family were Enterobacteriaceae (8.0% to 78%), followed by Enterococcaceae (1.7% to 64.2%) and Halomonadaceae (0.3% to 69.8%). The genera distribution showed great differences due to diet types and gut compartments. The fewest microbial species were shared by original populations, whereas the highest bacteria diversity was found for midgut of rice population feeding on water-oat. The bacterial communities in the midgut were more diverse than those in the hindgut. A comparison among phenotypically divergent populations of *C. suppressalis* shows that gut microbial communities vary with diet types and gut compartment.

## Background

The insects’ alimentary canal is a tube opening from the mouth to the anus, and is divided into three distinct regions, foregut, midgut and hindgut. Food is probably stored and partially digested in the foregut, fully digested and nutrients absorbed in the midgut, and useful materials and water are absorbed in the hindgut [1]. The alimentary canal is a desirable, nutrient-rich ecological niche where multiple microbial taxa flourish and reproduce. The anterior hindgut region is the most densely inhabited site of symbionts the alimentary canal, “due to the availability of partially digested food coming from the midgut, as well as the products excreted by the Malpighian tubules [2]”. The microbial taxa contribute to various functions, including nutrition, immune, development, survival, reproduction, detoxification [3–10] and population differentiation [11].

*Chilo suppressalis* is one of the destructive generalists of rice in Asia, southern Europe, and northern Africa [12–15]. The intercropping pattern (rice is planted in a mosaic fashion under a crop rotation system with water-oat) facilitates a transfer of *C. suppressalis* from rice plant to water-oat plant. After a long ecological adaptation, the *C. suppressalis* has diverged into phenotypically populations (i.e., rice population and water-oat population) [17–21]. The two populations exhibit significant phenotypic differences: morphometric differences [22–24], aliesterase isozymes and insecticide susceptibility [25], host preference [26, 27] and adaptability [28], biological characteristics [25, 26, 29–34](), supercooling points and glycerol content [35], photoperiodic response [36], genetic difference[37]) and transcriptomic difference [38].

It is suggested that the divergent population is related to microorganisms, and the symbionts are important factors promoting evolution of their insect hosts [11, 39]. To date, a few documents focused on the associations of insecticides and gut bacteria of C. suppressalis [40, 41]. However, whether the phenotypically divergent populations of *C. suppressalis* harbor different gut bacteria, less emphasis has been given. Herein, we characterize the microbial community structure of the water-oat and rice populations of *C. suppressalis* using next-generation sequencing.

## Methods and methods

### Specimen collection and rearing

Larval *C. suppressalis* of water-oat population were collected from the water-oat field in Lishui; rice population was collected from rice field in Yuyao, Zhejiang, China in 2016. Land owners of the two fields gave us permissions for sampling undertaken in this study. All larvae and plants were kept in an insectarium at 28±1°C, with a photoperiod of 16 h: 8 h (light/dark), and a relative humidity > 80%. For both populations, larvae were fed with water-oat fruit pulp and rice seedlings and the 4th instar larvae were sampled after three generations of rearing.

We analyzed the 16S rRNA gene to estimate the gut bacterial composition of the *C. suppressalis*: midgut and hindgut of water-oat population feeding on water-oat fruit pulp (JMG, JHG) and rice seedlings (jMG, jHG), midgut and hindgut of rice population feeding on rice seedlings (RMG, RHG) and water-oat fruit pulp (rMG, rHG), respectively. Both the water-oat and rice populations of *C. suppressalis* were fed with their original host until they molted into the adult form, and their eggs mass hatched into larvae. Then, they were separated from the colony, and reared on rice seedlings and water-oat, respectively. 150 individuals were analyzed for each diet condition.

### Experimental design

**Fig. 1.**
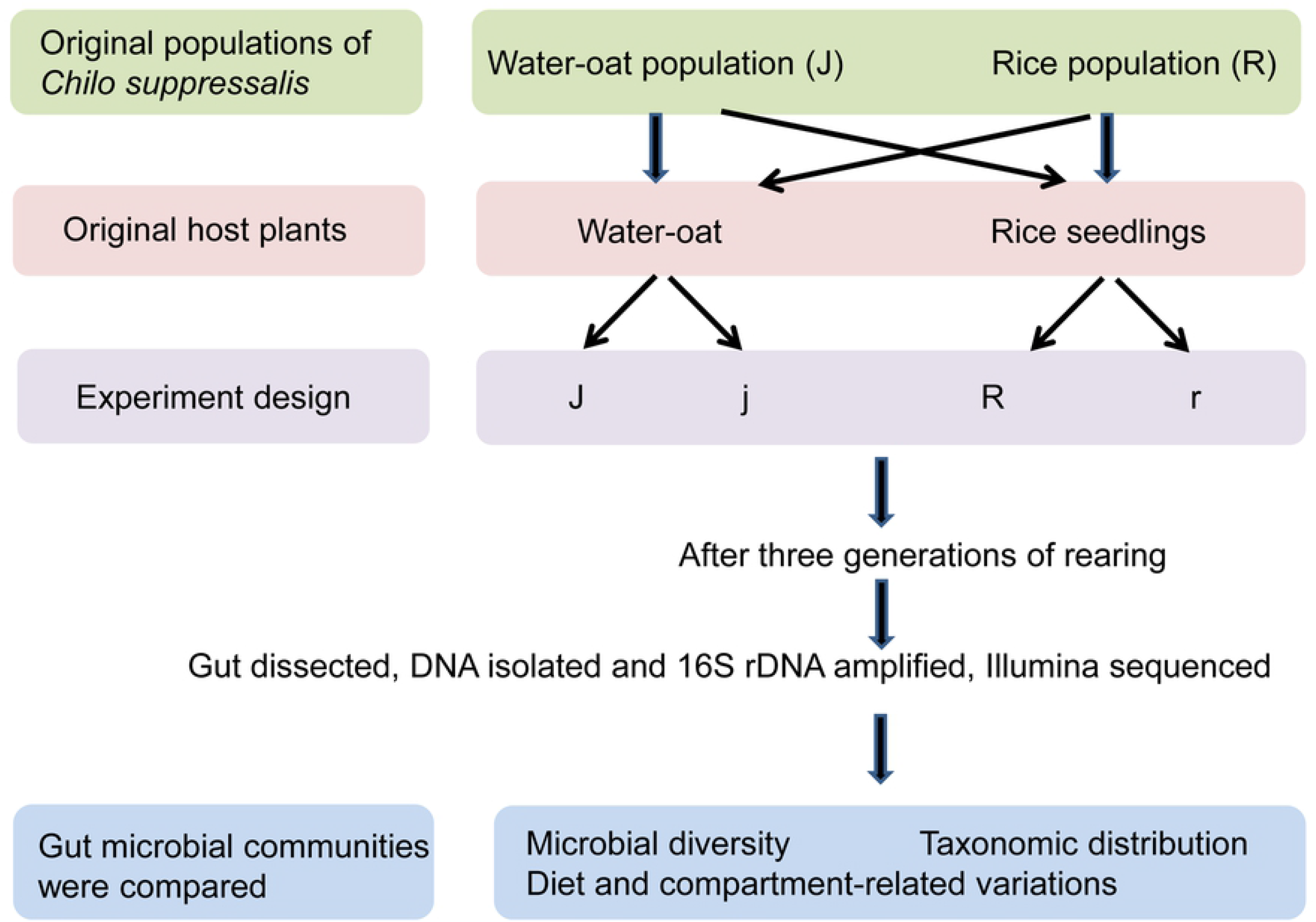
Schematic diagram of the fully factorial experimental design used in the present study. Chilo suppressalis from water-oat field were respectively reared on water-oat (J) and rice seedlings (j); and those from rice field were respectively reared on rice seedlings (R) and water-oat (r). All the groups were reared for three continuous generations to examine the effects of host plant, population origin and gut compartment on the gut microbial communities.

### *C. suppressalis* dissection and gut sample collection

Each individual was anesthetized by placed on ice and externally sterilized with 75 % and rinsed 3 times with sterile water. The gut were dissected out with sterilized fine-tip forceps and washed twice with sterile 0.9 % NaCl solution quickly. The midgut and hindgut were carefully separated and placed in different sterile microcentrifuge tubes. Each 50 midguts and hindguts were used as one sample, respectively. Three replicates were taken for each sample and immediately frozen in liquid nitrogen and stored at −80°C for DNA isolation.

### DNA isolation, 16S rDNA amplification

Total bacterial genomic DNA was extracted from the 8 sets of sample groups using a E.Z.N.A.^®^ Soil DNA Kit (Omega Bio-tek, Norcross, GA, U.S.) according to the manufacturer’s instructions. The DNA was finally eluted with TE buffer (Tris-EDTA buffer). DNA purity and concentration were measured using the NanoDrop 2000 spectrophotometer (Nano-drop Technologies, Wilmington, DE, USA). The total DNA was stored at −70°C until use.

The bacterial 16S rRNA variable V3-V4 regions were used to identify bacterial composition. Two universal primers (341F and 806R) which contain the specific barcode sequence were used for the amplification of the V4 region (341F: 5’-CCTAYGGGRBGCASCAG-3’, 806R: 5’-GGACTACNNGGGTATCTAAT-3’). The Polymerase Chain Reaction (PCR) reaction was performed in triplicate 20.0 μL mixture containing 4.0 μL 5×FastPfu Buffer, 2.0 μL 2.5 mM dNTPs, 0.8 μL of each Primer (5.0 μM), 0.4 μL FastPfu Polymerase, and 10 ng of template DNA. The amplification procedure was as follows: 95°C for 2 min, followed by 25 cycles of denaturation at 95°C for 30 s, annealing at 50°C for 30 s, and elongation at 72°C for 30 s and a final extension at 72°C for 5 min.

### Illumina MiSeq sequencing

Amplicons were extracted from 2% agarose gels and purified using a AxyPrep DNA Gel Extraction Kit (Axygen Biosciences, Union City, CA, U.S.) following to the manufacturer’s protocols and quantified using QuantiFluor™-ST (Promega, U.S.). Purified amplicons were pooled in equimolar and paired-end sequenced (2 × 250) on an Illumina MiSeq platform according to the standard instructions. The raw reads were deposited into the National Center for Biotechnology Information (NCBI) Sequence Read Archive (SRA) database (accession no. SRP116573).

### Processing of sequencing data

Raw fastq files were demultiplexed, quality-filtered using QIIME (version 1.17) with the following criteria: (i) The 250 bp reads were truncated at any site receiving an average quality score <20 over a 10 bp sliding window, discarding the truncated reads that were shorter than 50bp; (ii) exact barcode matching, 2 nucleotide mismatch in primer matching, reads containing ambiguous characters were removed; (iii) only sequences that overlap longer than 10 bp were assembled according to their overlap sequence. Reads which could not be assembled were discarded.

Operational Units (OTUs) were clustered using UPARSE (version 7.1 http://drive5.com/uparse/) and chimeric sequences were identified and removed using UCHIME. The phylogenetic affiliation of each 16S rRNA gene sequence was analyzed by RDP Classifier (http://rdp.cme.msu.edu/) against the silva (SSU115) 16S rRNA database.

## Results

### General structure of alimentary canal

The alimentary canal of *C. suppressalis* was a continuous tube running from the mouth to the anus. It was structurally divided into foregut, midgut, and hindgut. The foregut (Fg) is a slender, elongate tube, expanding posteriorly and constricts at its ends. The midgut (Mg) was a well-developed saclike tube beginning from the end of the foregut and extending to the long, narrow hindgut (Hg). In freshly dissected samples, the midgut is opaque white; the hindgut is yellowish-brown, whereas the foregut is translucent (Fig 2).

**Fig. 2.**
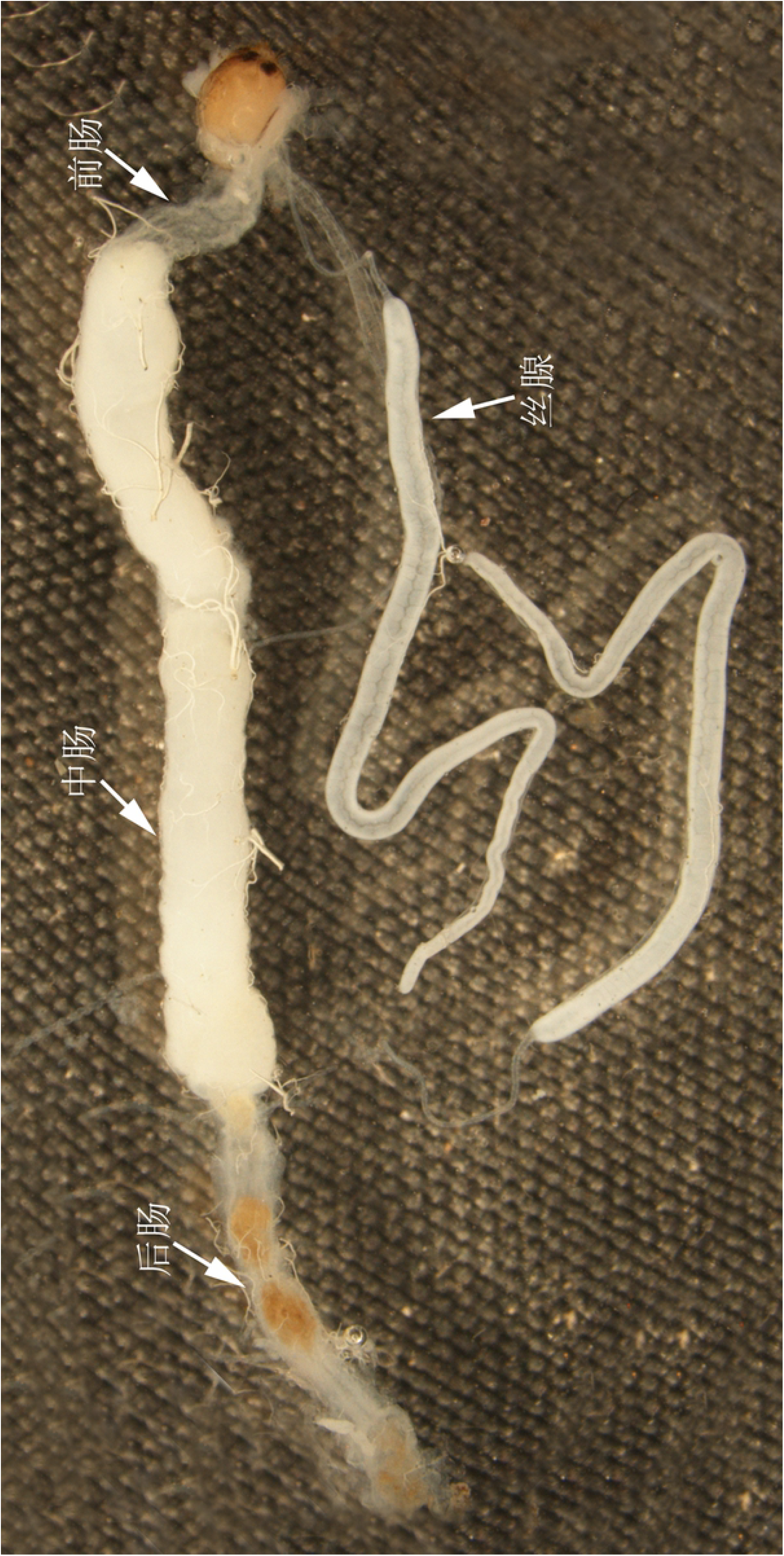
General structure of the alimentary canal of *Chilo suppressalis*. Fg, foregut; Mg, midgut; Hg, hindgut. Sg, salivary gland.

### Analysis of bacterial 16S rDNA gene sequences

Illumina sequencing obtained 861370 sequences that were clustered into 3234 OTUs (Table 1). Chao1 estimator and Shannon Index were calculated for the determination of the richness and homogeneity of the community. The relative bacterial abundance of 18 phyla differed significantly across eight samples (*Kruskal-Wallis* test, *p* < 0.0001). The midgut and hindgut of rice population feeding on water-oat (rMG and rHG) possessed the highest bacteria diversity, since their Shannon Index, Chao 1 estimator and total OTUs number were much higher. The bacteria of midgut samples were more diverse than those of hindgut samples (Table 1).

**Table 1.**
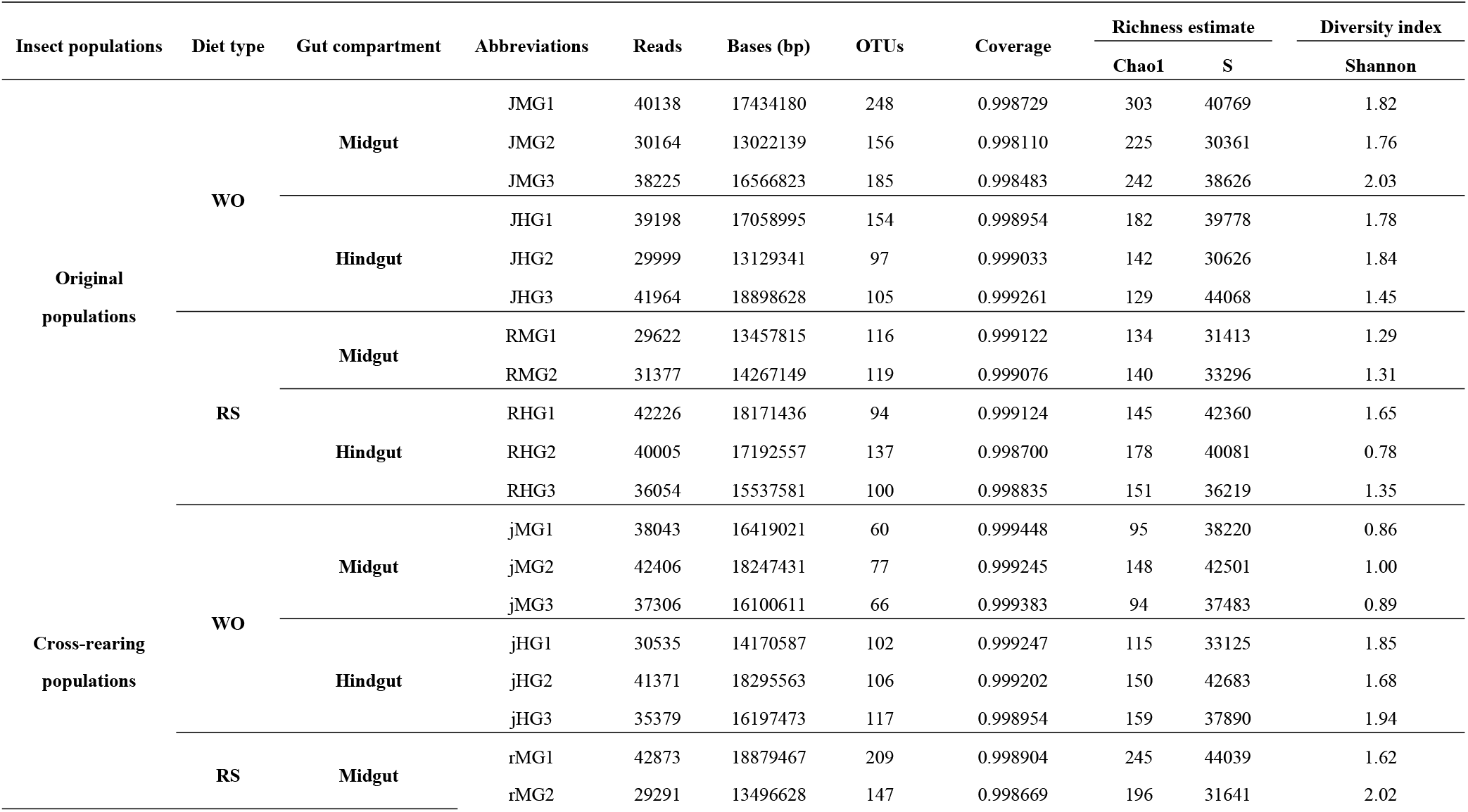
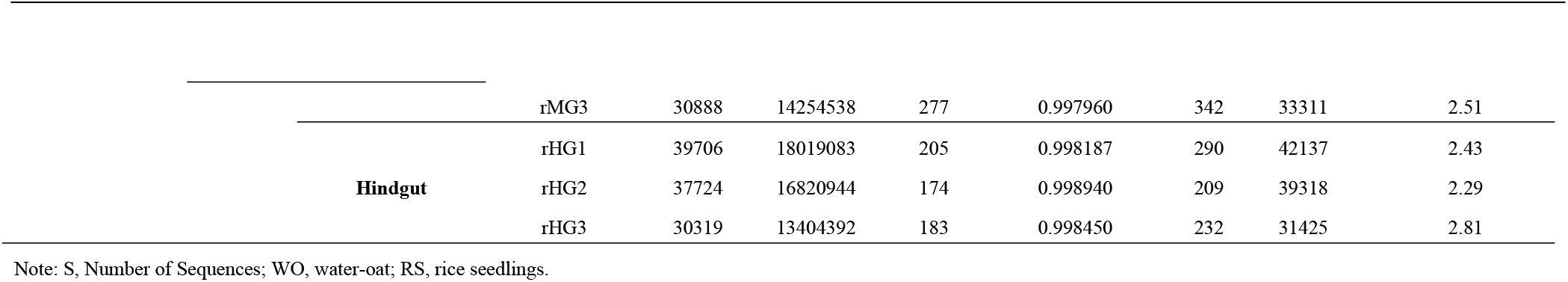
Diversity of gut bacterial communities based on sequencing.

### Microbial diversity of *C. suppressalis* gut microbiota

A total of 49 and 62 OTUs were observed in all midgut and hindgut samples respectively, indicating that a core set of species was prevalent in the bacterial communities of both compartments (Figs 3 and 4A, B). The core OTUs identified belongs to the phyla Proteobacteria, Firmicutes, Actinobacteria, Saccharibacteria and Bacteroidetes (S1 Fig). The OTUs were pooled into 31 core families for all midgut samples. The abundances of five families, i.e., Enterobacteriaceae (24.6%), Halomonadaceae (20.2%), Enterococcaceae (31.4%), Bacillaceae (11.4%), and Streptococcaceae (6.9%) (S1A Table; S2 Table and S3 Table). However, the OTUs were pooled into 28 core families for all hindgut samples. The abundant families were Enterobacteriaceae (66.4%), Enterococcaceae (11.2%), Bacillaceae (5.0%), Streptococcaceae (3.0%), Xanthomonadaceae (2.2%) and Flavobacteriaceae (1.7%) (S1B Table; S2 Table and S3 Table). rMG and rHG have the maximum number of unique OTUs, and the jMG and jHG possessed the minimum number.

**Fig. 3.**
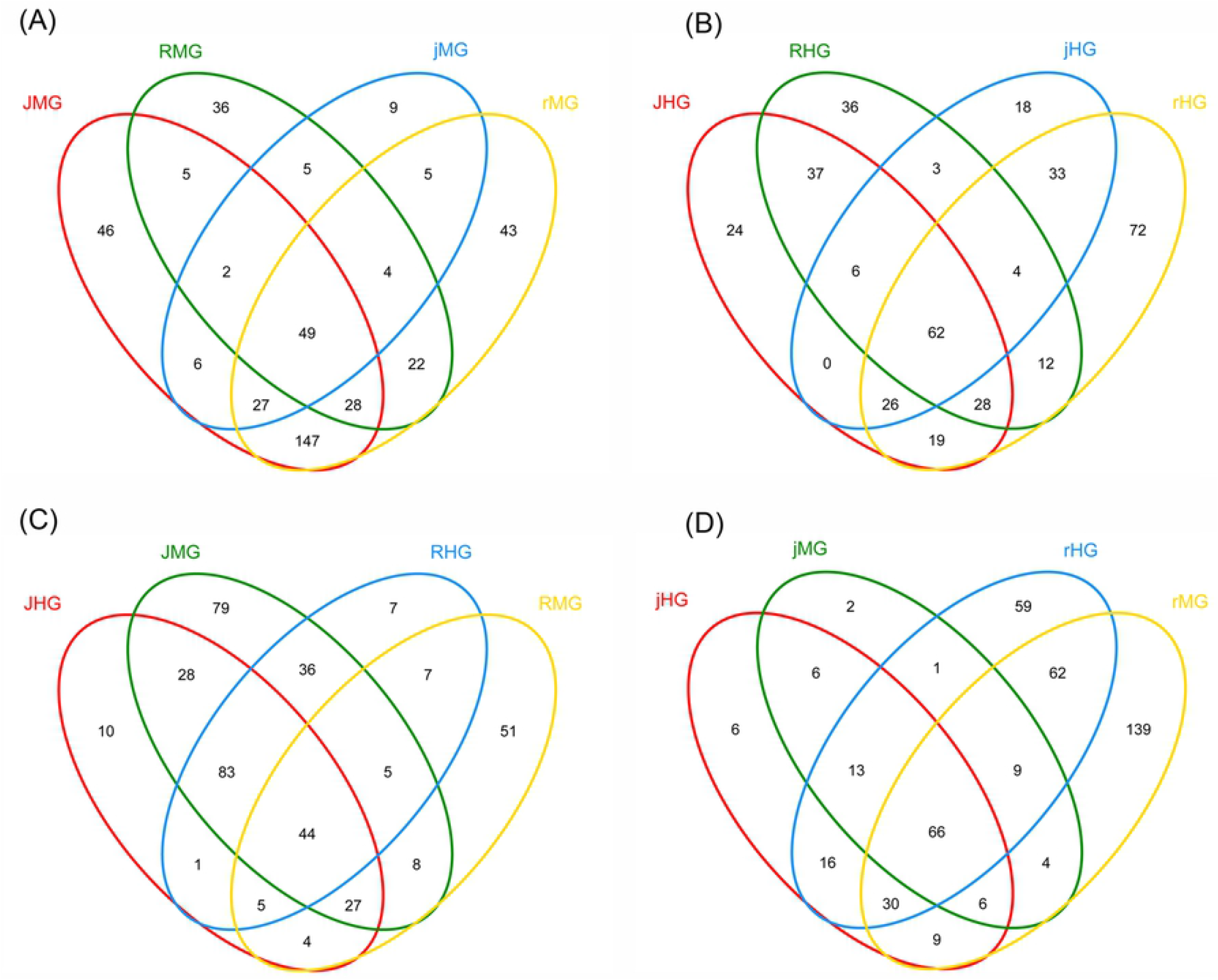
Core gut microbiota of *C.suppressalis*. Venn diagram representation of the OTUs from the different populations. Numbers inside indicate the OTUs shared by two or more samples, as well as unique families.

**Fig S1.** Bacterial composition (phylum level) of the microbiota along the midgut and hindgut of four different populations. (PDF 1688 kb)

To determine a core microbiota related to gut compartments changes in original populations and cross-rearing populations, we considered OTUs share from midguts and hindguts. A total of 44 and 66 OTUs were observed in gut of original populations and cross-rearing populations respectively (Fig 3C, D). The OTUs were pooled into 26 core families for midgut samples of original populations. The relative abundances of five families were Enterobacteriaceae, Halomonadaceae, Bacillaceae, Enterococcaceae and Streptococcaceae (S1C Table; S2 Table and S3 Table). However, the OTUs were pooled into 35 core families for hindgut of cross-rearing populations. Their abundant families were Enterobacteriaceae, Enterococcaceae, Streptococcaceae, Xanthomonadaceae and Halomonadaceae (S1D Table; S2 Table and S3 Table).

### Taxonomic distribution of insect gut bacteria

Taxonomic classification yielded 122 different families belonging to 18 different bacterial phyla (Fig 4), and the predominant phyla of all populations were Proteobacteria (16.0% to 96.4%), followed by Firmicutes (2.3% to 78.9%). At family level, Enterobacteriaceae (8.0% to 78%) was the most predominant taxa, and followed by Enterococcaceae (1.7% to 64.2%), and Halomonadaceae (0.3% to 69.8%) (S3 Table). There exhibit a high variation of relative abundance associated with diet and compartment, although the most abundant taxa were identified in all the samples.

**Fig. 4.**
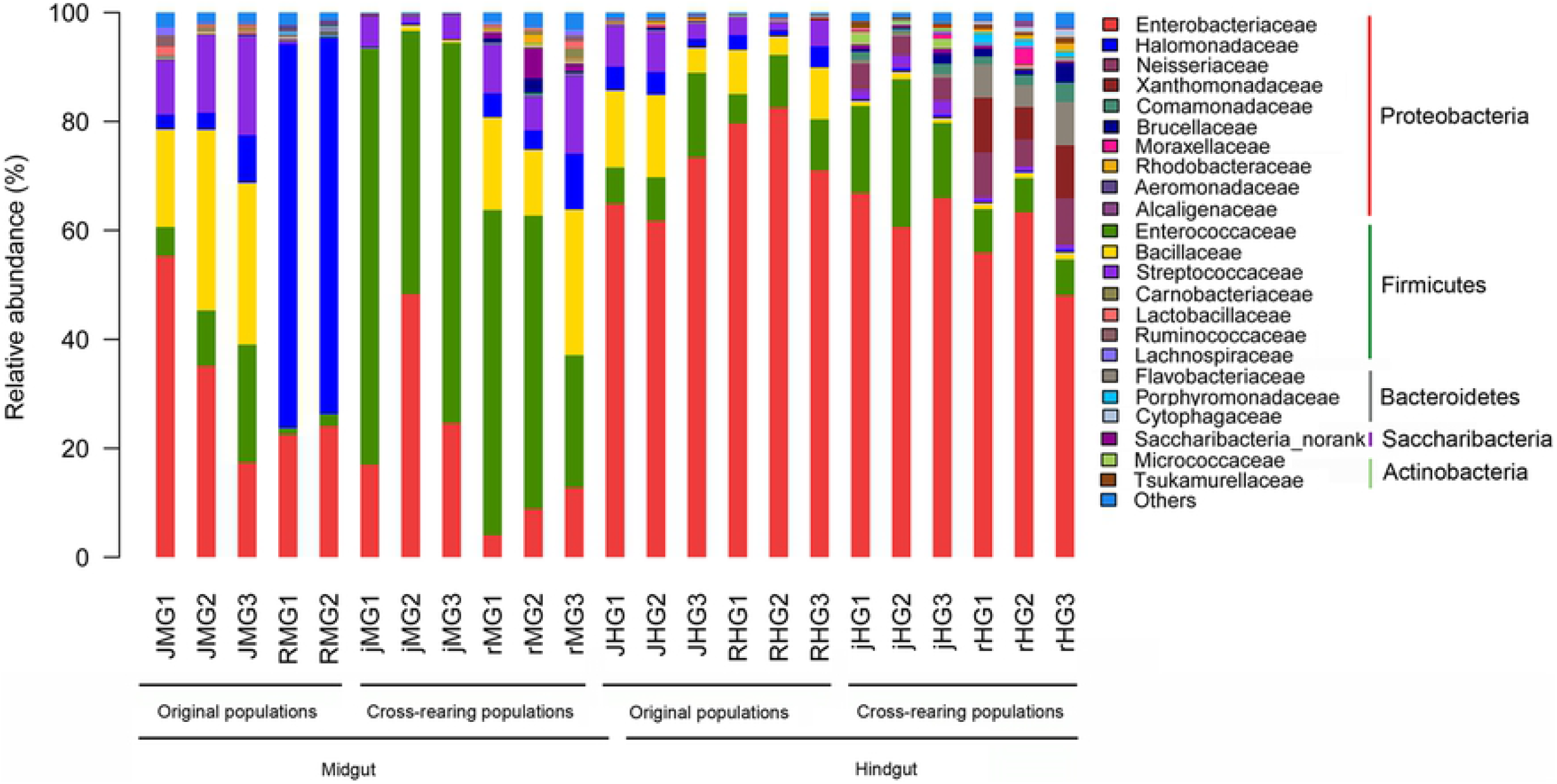
Bacterial composition (family level) along the midgut and hindgut of original and cross-rearing populations. Abbreviations for each sample are explained in Table 1.

Regardless of diet, a more homogeneous Phylum distribution was found for JHG, jHG, RHG and rHG: Proteobacteria (71.5% to 80.9%), Firmicutes (9.0% to 27.7%), Bacteroidetes (0.1% to 8.8%), Actinobacteria (0.1% to 2.8%) and Saccharibacteria (0.6%) respectively (Fig. 4; S1 Figure; S4 Table; S5 Table). However, the Firmicutes and Proteobacteria distribution changes separately occurred in JMG (40.3% to 71.8%, 27.3% to 58.6%), jMG (50.6% to 82.0%, 17.8% to 49.0%) and rMG (72.1% to 87.3%, 10.5% to 24.8%). Four bacterial phyla in RMG were more homogeneous in richness: Proteobacteria (96.2% to 96.6%), Firmicutes (2.2% to 2.3%), Bacteroidetes (0.6% to 0.9%) and Actinobacteria (0.3% to 0.5%).

**Fig. 5.**
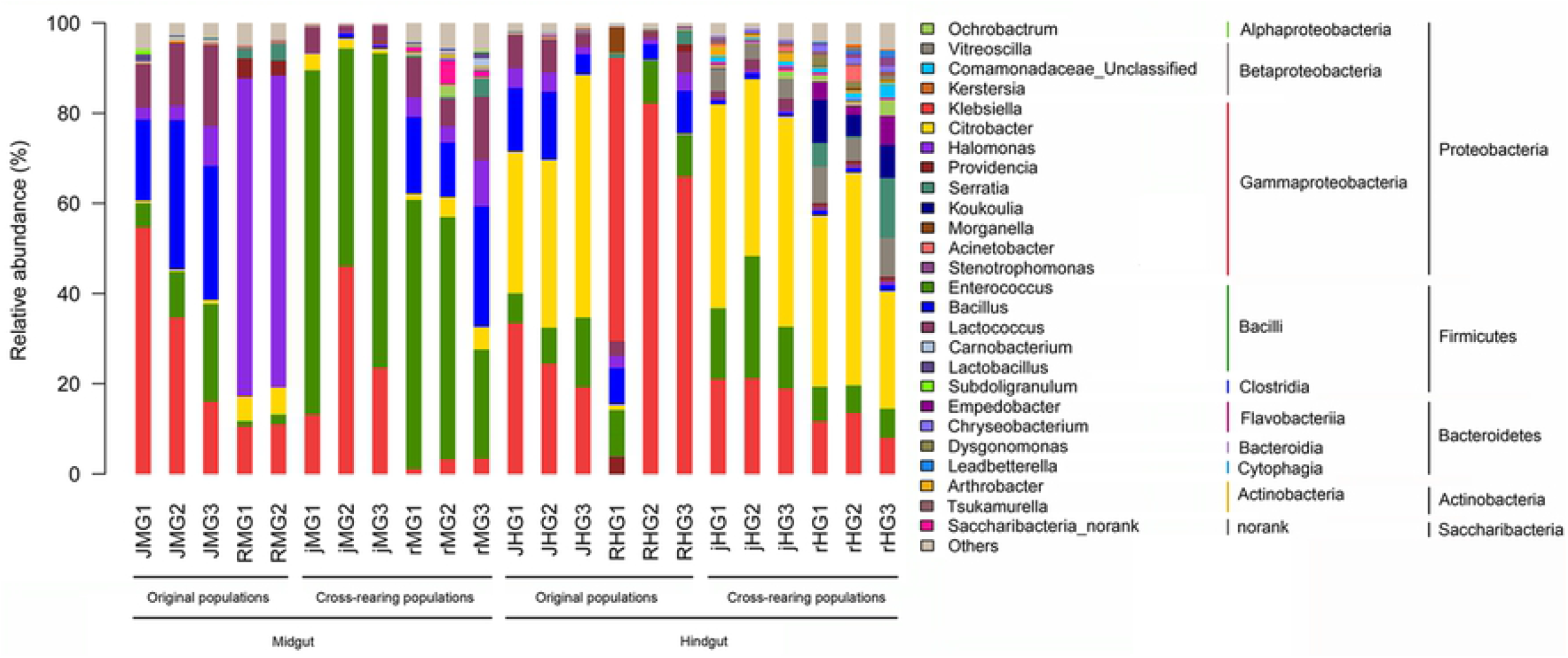
Bacterial composition (genus level) along the midgut and hindgut of original and cross-rearing populations. Abbreviations for each sample are explained in Table 1.

The bacterial genera from original populations showed distinct distribution according to diet types and gut compartments (Fig 5; S6 Table). *Halomonas* (69.9%) and *Klebsiella* (70.1%) were dominant in RMG and RHG, respectively; but *Bacillus* (26.9%) and *Klebsiella* (35.14%) were prevailed in JMG, *Citrobacter* (40.8%) was enriched in JHG. *Enterococcus* was dominant in jMG (64.8%) and rMG (45.9%), and *Citrobacter* was prevailed in jHG (43.7%) and rHG (37.1%). However, the bacteria in cross-rearing populations showed different genus distributions based on diet type. *Klebsiella* (27.6%) and *Bacillus* (18.7%) were the relative dominance in jMG and rMG, *Enterococcus* (18.9%, 6.7%) and *Klebsiella* (20.4%, 11.1%) were the relative prevalence in jHG and rHG, respectively.

### Diet and compartment-related variations in the microbial gut composition

There were significant differences in the relative abundances of microbial families in all gut samples (*p* < 0.0001, *Kruskal-Wallis* test). 95 bacterial taxa were identified at the genus level (Fig 6). Influence of compartment sampling proved significant with a well-defined cluster formed by JMG, jMG, rMG and RMG. By contrast, bacteria from RHG, JHG, jHG, rHG were more heterogeneous for constituting four different clusters. All the midguts and hindguts exhibit a significant difference in bacteria abundance of three main families: Enterobacteriaceae, Enterococcaceae and Bacillaceae. Enterobacteriaceae was dominant at hindgut (66.4%), but decreased to 24.6% at midgut. In comparison, Enterococcaceae was less abundant at hindgut (11.2%), while increased to 31.4% at midgut; Bacillaceae (5.0%) at hindgut, was increased to 11.4% at midgut (Fig 6; Table S4).

**Fig. 6.**
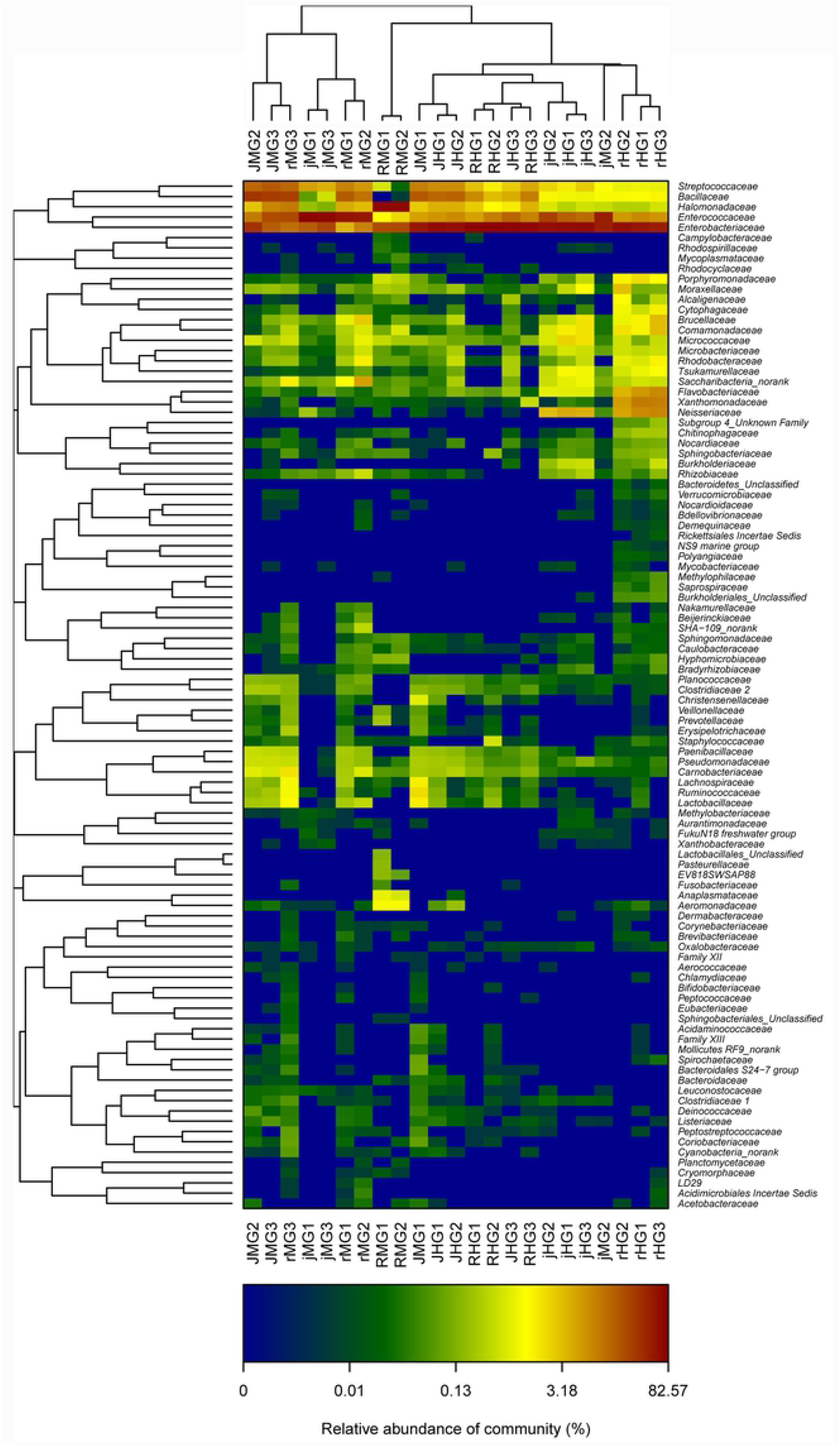
Heatmap and clustering of the midgut and hindgut microbiota of all populations. Heatmap colors show the percentage range of sequences assigned to the taxa.

Clustering was less consistent when its bacterial distribution is related to diet. jMG and rHG were the most homogeneous, which constituted an individual cluster; two of the jHG and rMG clustered together, but the other ones clustered with ones of the JHG and JMG, respectively. The rHG, jMG and RMG separately formed the most well-defined clusters; the rMG and JMG, jHG and JHG were more similar to each other. The difference at family level was a higher abundance of Enterobacteriaceae in the rHG (55.8%) than in rMG (8.6%) and JMG (35.9%); higher presence of Enterococcaceae in jMG (64.8%) than in rMG (45.9%), JMG (12.4%%), jHG (18.9%%) and JHG (10.0%); higher presence of Halomonadaceae in RMG (69.9%) than in rMG (6.0%), JMG (4.9%), jHG (0.3%) and JHG (3.4%). However, the Bacillaceae was higher in rMG (18.6%), JMG (26.9%) and JHG (11.3%) than in rHG (1.2%) and jMG (0.6%) (Fig 6).

A non-metric multidimensional scaling (NMDS) analysis was performed for analyzing influence of diet and compartment on the microbiota (Fig 7A-D). At midgut, the clusters were well defined and the highest variability was found in the RMG cluster. The RMG and jMG clusters exhibited the most different taxa composition, followed by the rMG and JMG clusters, showing an intermediate composition (Fig 7A). At hindgut, there were clearly separated clusters: the RHG clusters exhibited a higher intersample variation; the JHG cluster showed an intermediate composition respect to the RHG, jHG and rHG clusters (Fig 7B). JMG, JHG, RMG, RHG clusters were well-defined, and the JMG, RHG clusters had similar homogeneity level (Fig 7C). RMG was the most heterogeneous, followed by JMG and RHG. Clusters of cross-rearing populations were better defined than those of original populations. rMG was the most heterogeneous in taxa composition, followed by the jHG.

**Fig. 7.**
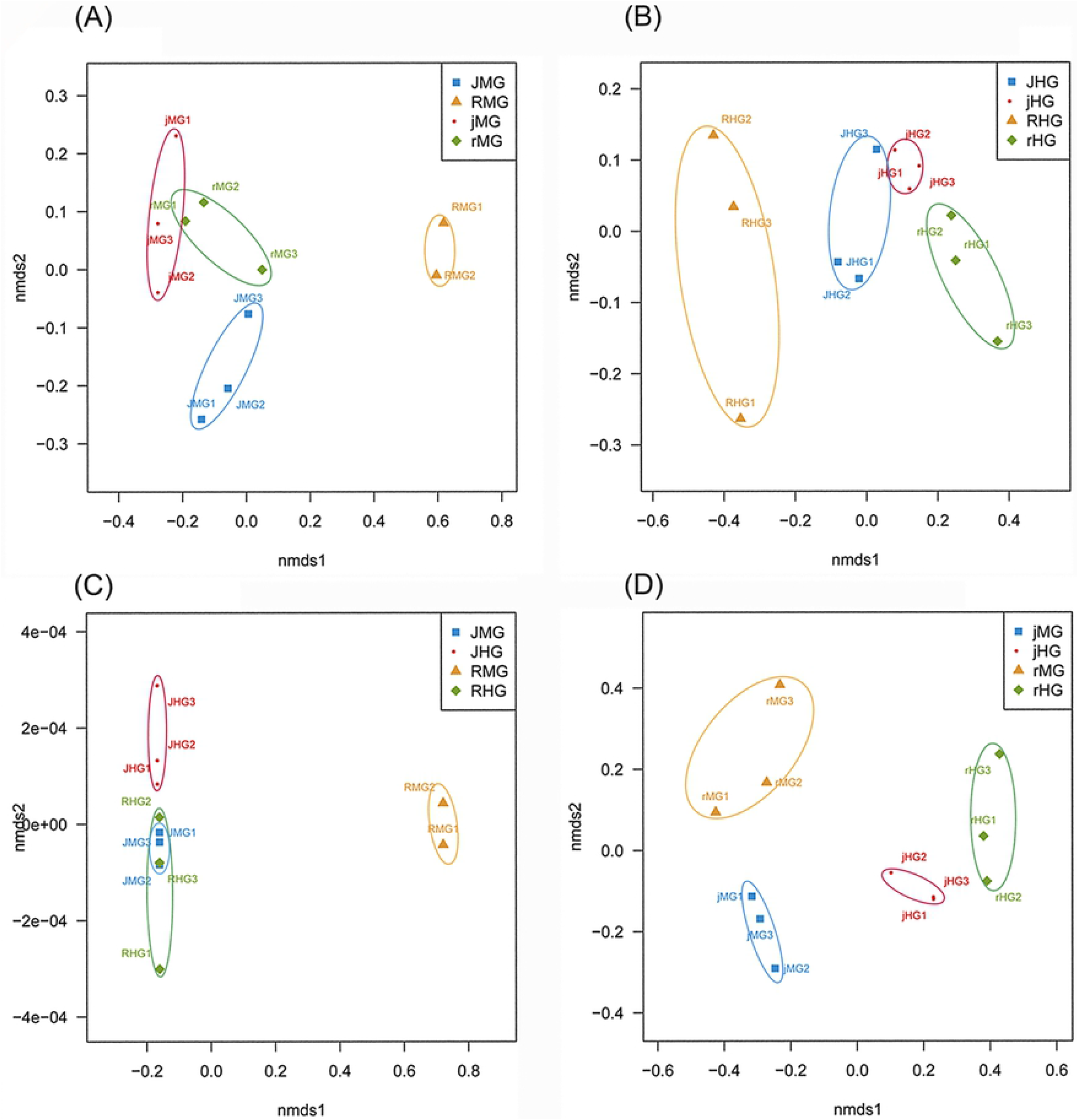
NMDS of the *C. suppressalis* gut microbiota. (A) NMDS of the taxon distribution of midgut samples. The samples were clustered by diets and represented with different colors: jMG (red, circles), rMG (green, rhombus), JMG (blue, squares) and RMG (saffron yellow, triangle). (B) NMDS of the taxon distribution of hindgut samples. The samples were clustered by diets and represented with different colors: jHG (red, circles), rHG (green, rhombus), JHG (blue, squares) and RHG (saffron yellow, triangle). (C) NMDS of the taxon distribution of midgut and hindgut samples from water-oat and rice populations. The samples were clustered by diets and represented with different colors: JHG (red, circles), RHG (green, rhombus), JMG (blue, squares) and RMG (saffron yellow, triangle). (D) NMDS of the taxon distribution of midgut and hindgut samples from cross-rearing populations. The samples were clustered by diets and represented with different colors: jHG (red, circles), rHG (green, rhombus), jMG (blue, squares) and rMG (saffron yellow, triangle). The ellipses represent the standard error of the centroid for each group of samples with a confident limit of 95%.

## Discussion

### Bacterial diversity and distribution

To date, there are few documents on how gut microbial communities differ across divergent insect populations based on diet and gut compartments. Gut bacterial diversity overall was notably greater in water-oat-fed rice population of *C. suppressalis* compared to rice-fed one or water-oat populations; midgut bacteria were more diverse and variational than hindgut ones. Only bacteria of *Citrobacter, Enterococcus, Halomonas*, and *Klebsiella* were shared by original populations. The comparative distribution in *C. suppressalis* suggests that they are core gut microbiota, and it is probable that these microbes are beneficial to their hosts. Rice seedlings and water-oat are very different in nutritional ingredient and secondary compounds would have different microbes available.

The gut bacterial composition and richness exhibited significant differences in the midgut and hindgut of different populations of *C. suppressalis. Halomonas* and *Klebsiella* dominated the midgut and hindgut of rice-fed rice population, and *Klebsiella* and *Citrobacter* are prevailed the midgut and hindgut of water-oat-fed water-oat population. *Enterococcus* dominated the midgut of cross-rearing populations, and *Citrobacter* was found exclusively in the hindgut of cross-rearing populations. The genus is characterized by the ability to convert ethanol to acetic acid in the presence of oxygen [42], and it has not been reported previously in rice stem borers. *Enterococcus* is associated with insecticide and pathogen resistances [40, 43], and the presence of this genus in the *C. suppressalis* suggests the enhancement of this pest’s immune system during host shift.

A gut ‘microbial core’ appears to be common, though an inter-individual variability exists in *C. suppressalis*. The inter-individual variability was previously documented in honey bees *Apis mellifera* [44], Anopheles [45], and cockroaches *Blattella germanica* [46, 47], *Shelfordella lateralis* [48] and *Periplaneta americana* [49]. Curtis and Sloan (2004) suggested that the variation could be attributed to the “random acquisition of microorganisms from a highly diverse environmental reservoir community” [50]. However, as the populations were reared many generations in laboratory with identical conditions, such variability could be attributed to host genetics and population divergence. Cluster analysis showed the jHG and rHG formed the most well-defined clusters and suggested stable microbial profiles, whereas the rMG and JMG, JMG and JHG were more similar to each other. The inter-individual differences suggested that SSB gut microbiome profiles may serve as useful biomarkers for bio-control in population-based studies.

### The effect of diet on gut microbial composition

We found two dominant phyla (i.e., Proteobacteria and Firmicutes) and three families are differed significantly in abundance, indicating that a rapid fluctuation may be related to the adaptation for new host or environments. Proteobacteria were reported to be involved in carbohydrate degradation, such as starches and hemicellulose [51], and can be involved in pectin-degrading [52] and nitrogen [53]. The oligophagous diet of stem borers provides suitable ecological niches for harboring bacteria in compared with monophagous lepidopterans [54]. Comparison of different diets indicated that the diet is an important factor in modulating the structure of bacteria community among populations of *C. suppressalis*, as was documented for other insect species [46, 54–58].

The gut bacterial genera are also varied, due to the difference of diets in *C. suppressalis*: in original populations, *Halomonas* was dominant in the RMG, *Klebsiella* was prevailed in RHG and JMG, and *Citrobacter* was dominant in the JHG; in cross-rearing populations, *Enterococcus* was dominant in midgut, and *Citrobacter* was prevailed in hindgut. Since diet and host taxonomy structure bacterial microbiome composition [55, 59], the successful expansion of bacteria over time probably in turn suppressed the growth of bacteria from other phyla in the same habitat [43]. We infer that the different bacteria dominance might be related to successful reproduction of some bacteria genus and suppression of the other ones.

Although water-oat and rice plants are phylogenetically close and belonged to the tribe *Oryzeae*, better biological performances (including better growth and survival rate) of *C. suppressalis* fed on water-oat than on rice reflect great differences in the nutriments and secondary substances between the two host plants [21, 32, 60–61]. Members of Xanthomonadaceae exhibits cellulase activity [60], and also play a role in the metabolism of lignin-derived aromatic compounds [62]. Members of *Halomonas* were reported to have cellulolytic activity [63, 64]. Thus colonize of these bacteria indicate the degradation of nutrient component appears to be important for *C. suppressalis* during its feeding on rice seedlings or water-oats. In addition to differences in nutritional quality, the water-oat and rice seedlings possess different allelochemicals: the water-oat contains caffeic acid, gallic acid and cinnamic acid [65]; whereas the latter has oxalic acid, total phenolic and tannin [66]. Dietary lignocellulose composition could cause shifting rapidly in the gut microbiota [58]. The gut bacteria of *C. suppressalis* may change allelochemicals come from water-oat and rice seedlings via complex process.

One interesting and unexpected result concerns the two compartments chosen for analysis, as we found that variability in microbial composition is higher in midgut than in hindgut, independently of diet. The obvious community difference indicates that only some specific groups of microorganisms are able to survive and colonize in the hindgut. The comparison of the microbiota composition of midgut and hindgut of *C. suppressalia* fed the same diet provide insights into the compartment changes in the gut microbiota of SSB. Since both gut regions are alkaline in *C. suppressalis*, it is likely that factors other than pH are responsible for this shift. Rather, the force driving community structure could be the unique morphology, favorable physiological conditions (*viz*., oxygen content), lack of various enzymes and the availability of partially digested food lead the hindgut of *C. suppressalis* to become a benign site for maintaining special bacteria.

## Conclusions

In this study, we have investigated the gut microbial communities of phenotypically divergent populations in *C. suppressalis* after feeding on original and non-original hosts. The results showed that the highest bacteria diversity was found for midgut of rice population feeding on water-oat. The most dominant phyla were Proteobacteria and Firmicutes; and the dominant families were Enterobacteriaceae, followed by Enterococcaceae and Halomonadaceae. The microbial communities are highly diverse at the genera level due to diet types or gut compartments among populations. The bacterial community composition is driven mainly by diet types, and affected by other factors including gut compartments. These findings provide an important insight into investigation of insect-bacteria symbioses, and biocontrol of this species and other lepidopterans.

## Abbreviations

SSB: Striped stem borer;
NMDS: Non-metric multidimensional scaling;
PCR: Polymerase chain reaction;
OTUs: Operational Units;
JMG: Midgut of water-oat population;
JHG: Hindgut of water-oat population;
RMG: Midgut of rice population;
RHG: Hindgut of rice population;
jMG: Midgut of water-oat population feeding on rice seedlings;
jHG: Hindgut of water-oat population feeding on rice seedlings;
rMG: Midgut of rice population feeding on water-oat;
rHG: Hindgut of rice population feeding on water-oat;
NCBI: National Center for Biotechnology Information;
SRA: Sequence Read Archive

## Acknowledgements

The authors are grateful to Pan Xiaoting for rearing the striped stem borer.

## Authors’ contributions

**Experiments conceive and design**: Jianming Chen, Haiying Zhong.

**Experiments performance, data analysis, and result evaluation:** Haiying Zhong.

**Writing – original draft:** Haiying Zhong.

R**esults discussion and manuscript revision:** Jianming Chen, Haiying Zhong.

**Collecting and rearing insect populations:** Juefeng Zhang, Fang Li.

## Availability of data and materials

The sequence data used in this study was deposited into the National Center for Biotechnology Information (NCBI) Sequence Read Archive (SRA) database (accession no. SRP116573).

## Supporting information

**S1 Fig.** Bacterial composition (phylum level) of the microbiota along the midgut and hindgut of four different populations. (PDF 1688 kb)

**S1A Table.** OUT name of bacteria of all midgut samples in venn. (PDF 46 kb)

**S1B Table.** OUT name of bacteria of all hindgut samples in venn. (PDF 45 kb)

**S1C Table.** OUT name of bacteria of midgut and hindgut samples of two original populations in venn. (PDF 46 kb)

**S1D Table.** OUT name of bacteria of midgut and hindgut samples of two cross-rearing populations in venn. (PDF 47 kb)

**S2 Table.** Classification table of bacteria species. (PDF 204 kb)

**S3 Table.** Mcrobial community percent at family level (merged biological replicates). (PDF 65 kb)

**S4 Table.** Mcrobial community percent at family level (unmerged biological replicates). (PDF 80 kb)

**S5 Table.** Mcrobial community percent at phylum level. (PDF 40 kb)

**S6 Table.** Mcrobial community percent at genus level. (PDF 177 kb)

